# eMIND: Enabling automatic collection of protein variation impacts in Alzheimer’s disease from the literature

**DOI:** 10.1101/2023.09.07.556602

**Authors:** Samir Gupta, Xihan Qin, Qinghua Wang, Julie Cowart, Hongzhan Huang, Cathy H Wu, K Vijay-Shanker, Cecilia N Arighi

## Abstract

Alzheimer’s disease and related dementias (AD/ADRDs) are among the most common forms of dementia, and yet no effective treatments have been developed. To gain insight into the disease mechanism, capturing the connection of genetic variations to their impacts, at the disease and molecular levels, is essential. The scientific literature continues to be a main source for reporting experimental information about the impact of variants. Thus, development of automatic methods to identify publications and extract the information from the unstructured text would facilitate collecting and organizing information for reuse. We developed eMIND, a deep learning-based text mining system that supports the automatic extraction of annotations of variants and their impacts in AD/ADRDs. In particular, we use this method to capture the impacts of protein-coding variants affecting a selected set of protein properties, such as protein activity/function, structure and post-translational modifications. We conducted an evaluation on the efficacy of eMIND to extract variant impact relations and obtained a recall of 0.84 and a precision of 0.94. The publications and extracted information are integrated into the UniProtKB computationally mapped bibliography to expand annotations on protein entries. eMIND’s text-mined output are presented using controlled vocabularies and ontologies for variant, disease and impact along with the evidence sentences. A sample of annotated abstracts can be accessed at URL: https://research.bioinformatics.udel.edu/itextmine/emind.

## 1 Introduction

Alzheimer’s Disease and related dementias (AD/ADRDs, e.g., Frontotemporal dementia, Lewis body dementia, and Nasu-Hakola Disease) are multifactorial disorders and constitute the most common forms of dementia. Over 55 million people are currently living with dementia, with 60 to 70% of the cases corresponding to Alzheimer’s disease (World Health Organization). With nearly 10 million new cases every year and no effective interventions, it is critical that the data generated by the AD/ADRD research is organized and available in data resources to increase the pace of discovery and innovation. Specifically, to understand the underlying disease mechanisms, it is highly relevant to collect of candidate genes, their variants and the impact of such variants at the molecular level.

UniProt (UniProt Consortium, 2021)(Universal Protein Resource, RRID:SCR_002380), a publicly available and comprehensive resource of protein sequence and function information, has been actively annotating Alzheimer’s relevant proteins in coordination with the AD expert community (Breuza et al., 2020) in its knowledgebase (UniProtKB). In the context of expert annotation of natural variants, priority is given to those that are linked to disease and have impact on some property at the molecular level. For example, the variant p.Gly590Arg of tau (MAPT, UniProtKB:P10636) is linked to Frontotemporal dementia, and this variant impacts the aggregation propensity of tau and binding affinity of tau towards microtubules and F-actin (Sandberg et al., 2020). With the aim of broadening the spectrum of publications associated with a protein entry, UniProt collects additional publications and annotations from external resources (e.g., Alzheimer’s Research Forum, RRID:SCR_006416) and also contributed by the community (Wang et al., 2021). In addition, large-scale variant data is imported from a variety of resources (e.g., 1000Genomes, RRID:SCR_006828; ClinVar, RRID:SCR_006169; COSMIC, RRID:SCR_002260; Exome Sequencing Project, RRID:SCR_012761; ExAc, RRID:SCR_004068) to complement the set of variants captured from the literature. The scientific literature remains a main source where experimental data about variants and their impact are described. Thus, here we present eMIND, a text mining system that supports the identification of publications and extraction of impact annotations in AD/ADRDs for integration into the UniProtKB computationally mapped bibliography.

In particular, the impact types captured by eMIND, shown in this work, are those that are most relevant to UniProt for variant effect annotations (**Figure 1, molecular level**), including impact of variation on protein activity, interaction, post-translational modification, structure, biological process, processing, protein abundance, and protein aggregation. The last three are of special importance in the context of AD/ADRDs, as these are predominantly mentioned in these disorders. Although, eMIND is capable of capturing impact at disease level (**Figure 1**), e.g., risk to disease, phenotype, and disease onset, its evaluation is ongoing and out of the scope of this work.

**Figure 1:**
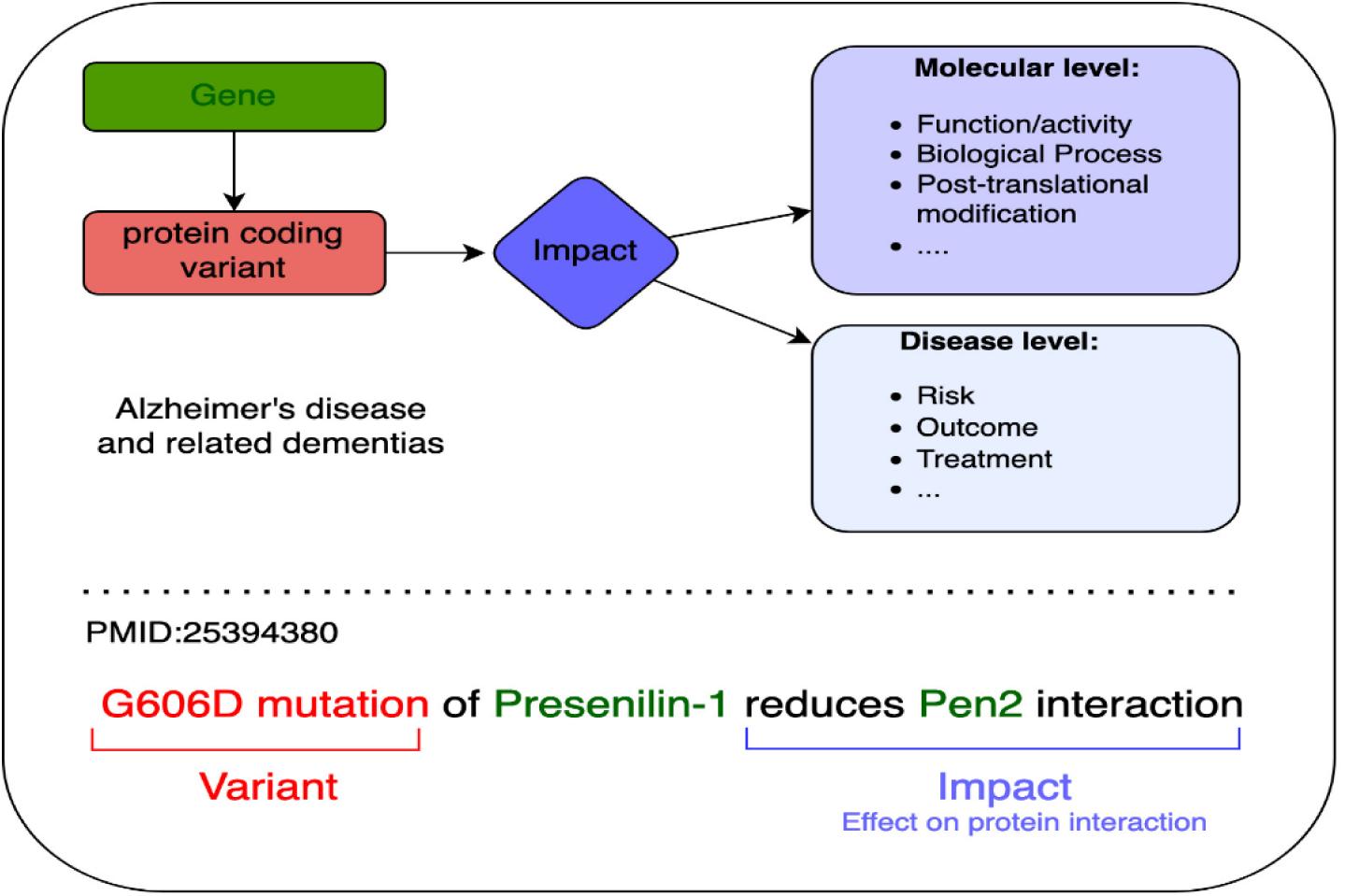
Illustration of eMIND system, capturing gene variants and their impact in the context of AD/ADRDs. The effect of a variation is divided into two levels, namely molecular level and disease level. Example sentence highlighting variant (red), genes (green) and a molecular level impact (blue) for a molecular level impact.

eMIND approach involves recognition of different types of impacted entities, detection of phrases describing genetic variants (mutations) and a relation extraction classifier to extract impact relations between the variation and the impacted entities within a sentence. The relation extraction classifier is built using transformers, specifically the widely popular pre-trained BERT model (Devlin et al., 2019). We conducted an evaluation on the efficacy of eMIND to extract variant impact relations and obtained a recall of 0.84 and a precision of 0.94.

UniProt is working towards structuring the description of the functional effects of amino acid modifications, including those of variants. For this, it will use terms in the variation ontology VariO (Vihinen, 2014) (Variation Ontology, RRID:SCR_000311) to describe the effect of a variant in combination with a target when needed. For instance, the UniProt variant annotation for p.Gly590Arg of tau (MAPT, UniProtKB:P10636) mentioned before could be structured using the following terms:VariO:0038 effect on protein aggregation and VariO:0058 effect on protein interaction, with target: microtubules and F-actin. With this into consideration, annotations in eMIND are presented using VariO terms for the impact types, along with other controlled vocabularies, such as HGVS for variant nomenclature (den Dunnen et al., 2016) (HGVS Locus Specific Mutation Databases, RRID:SCR_006730), disease ontology (Schriml et al., 2022) (Disease Ontology, RRID:SCR_003491), gene ontology (GO Consortium, 2021) (Gene Ontology, RRID:SCR_002811) and VariO, in addition to the evidence sentence. eMIND output on a sample set of AD/ADRDs related abstracts is available online at URL:https://research.bioinformatics.udel.edu/itextmine/emind and can be downloaded in JSON format.

## 2 Background and Related Work

### 2.1 Mutation Impact Types

The mutation impact types captured by eMIND include impact at the molecular level relevant for UniProt such as impact of variation on protein activity, interaction, post-translational modification, structure, biological process, processing, protein abundance, and protein aggregation (Table 1). Additionally, eMIND also considers variant impact at the disease level i.e. impact in the context of AD/ADRD such as risk to disease, phenotype, and disease onset (Table 1, last three rows), its evaluation is ongoing and out of the scope of this work.

**Table 1:** Mutation impact types.

| Impact Type | Example Sentence |
| --- | --- |
| **Protein Processing** | Arctic mutation favors proamyloidogenic <u>APP processing</u> by increased <u>beta-secretase cleavage</u>...” |
| **Protein Interaction** | ..Alzheimer's disease-protective 'Icelandic' mutation greatly attenuates <u>APP-BACE-1 interactions</u>, suggesting a mechanistic basis for protection. |
| **Protein Abundance** | <u>Cerebrospinal fluid amyloid-beta levels</u> were influenced by PPAR-alpha L162V genotype. |
| **Protein Post-Translational Modification** | A single Y202F mutation in the Mint1 N-terminus inhibits C-Src binding and <u>tyrosine phosphorylation</u>. |
| **Protein Activity** | ...the expression of the N141I PS2 FAD gene strongly promoted (>2-fold) <u>glycogen synthase kinase (GSK)-3beta activity</u> .. |
| **Protein Aggregation** | Arctic APP mutation is sufficient to cause <u>amyloid deposition</u> .. |
| **Protein Structure** | The L392V mutation of presenilin 1 associated with autosomal dominant early-onset Alzheimer's disease alters the <u>secondary structure of the hydrophilic loop</u>. |
| **Biological Process** | ...cell cycle progression was more inhibited by the expression of PS2(N141I) compared with wild-type PS2. |
| **Disease Risk** | .S311C polymorphism (SS genotype) and the rs705379 (the A allele and GG genotype) are associated with <u>risk of AD</u>.. |
| **Disease Phenotype** | These findings expand the number of APP mutations linked to <u>hereditary cerebral hemorrhage with amyloidosis</u>, reinforcing the link between this phenotype and codon 693 of APP. |
| **Disease Onset/Pathogenesis** | The K311R mutation might contribute to <u>AD pathogenesis</u>.. |

### 2.2 Impact Constructs

Our notion of impact of an entity was motivated by our previous work on miRiaD (Gupta et al., 2016), a tool that can extract associations between microRNAs and diseases from the literature. During the development of miRiaD, we noticed that nearly all statements connecting microRNAs to biological processes and disease-related terms were through mentions of any of the following types of relation: regulation (e.g., miR-31 “regulates/promotes/inhibits” apoptosis), involvement (e.g., miR-31 “is involved in/plays a role in” apoptosis) and association (miR-31 “was associated with/was correlated with” apoptosis). We termed this set of relations as CAIR relations (Connections through Association, Involvement, and Regulation). It soon became evident that CAIR relations are not specific to microRNAs and are generalizable to other entity types such as genes, proteins, and variants. Thus, we generalized the concept of the **impact** of entities (“variant/mutation”) and events (e.g., “Phosphorylation of protein”) by developing a relation extraction (RE) system to extract such CAIR relations based on the hypothesis that such relations are widely used in connecting biomolecular entities with diseases, processes and other events, which are important for capturing effect/impact of entities to its environment. Note, the notion of impact through CAIR relations is independent of the concept types it connects, and thus, it can be extended to extract impact between different biomedical entities and processes. In this work, we use the CAIR relations extracted by the rule-based miRiaD system to train a BERT-based mutation impact extraction model. Examples sentences of the impact relation constructs we consider in eMIND are shown in Table 2.

**Table 2:** Impact relation constructs.

| **Relation Construct** | **Example Sentence** |
| --- | --- |
| **Regulation** | G206D mutation of presenilin-1 <u>reduces</u> Pen2 interaction. |
| **Involvement** | <u>Involvement</u> of paraoxonase 1 genetic variants in Alzheimer's disease neuropathology. |
| **Association** | S311C polymorphism (SS genotype) and the rs705379 (the A allele and GG genotype) are <u>associated with</u> risk of AD. |

### 2.3 Related Works

Few corpora and annotation guidelines have been developed and reported, which aim to capture certain causal relations between biomedical entities and events. An annotation scheme and corpus containing causal biomedical event-level relations were described in BioCause (Mihăilă et al., 2013). Even though the notion of causality described in BioCause closely relates to our notion of impact (through CAIR relations), BioCause focuses on causal associations in discourse (coherent sequence of clauses and sentences) rather than associations between events or entities. In addition, we wish to note the existence of a corpus for the detection of bio-molecular events in the BioNLP Genia Event extraction Shared task 2011 (Kim et al., 2011). The GENIA Event Corpus and works developed based on it are limited to regulation events between a small set of entities: protein, an event involving a protein (expression, phosphorylation, etc.), and site.

There has been some work on extracting broad biomedical semantic and causal relationships between entities and processes from the biomedical text, which involve generally rule-based methods. SemRep (Kilicoglu et al., 2020) employs a pattern-based approach for identifying relationships between a pair of entities. However, these entities and relationships are limited to the UMLS Metathesaurus and Semantic Network and do not cover the range of impact that we considered in the CAIR relations. BELMiner (Ravikumar et al., 2017) is a rule-based RE system to extract events in biological expression language (BEL). The BEL framework was developed as part of the BioCreative V Track4 challenge allowing for a formal representation of causal relationships between BEL terms, which were limited to certain findings or observations (sample abundance and processes).

Additionally, several text-mining applications have been developed for specific biomedical RE tasks, which include relations and events relevant to this work. Among these, protein-protein interactions (Tikk et al., 2010; Peng et al., 2015; Peng and Lu), post-translational modifications (Xu et al., 2012; Torii et al., 2014), mutation-gene associations (Mahmood et al., 2016; Allot et al., 2018), subcellular localization (Zheng and Blake, 2015; Cejuela et al., 2018; Su et al., 2019) and chemical protein interactions (Peng et al., 2018; Antunes and Matos, 2019). Earlier approaches used rule-based methods, but due to the growing popularity of deep learning approaches, recent works have focused on deep learning architectures, such as Convolutional Neural Networks (CNN) and Long short-term memory (LSTM). Lately, the advent of transformer-based models, such as BERT (and its variants, such as BioBERT) have achieved remarkable success in various RE tasks as noted in the Biomedical Language Understanding Evaluation (BLUE) benchmark (Peng et al., 2019) and Biomedical Language Understanding and Reasoning Benchmark (BLURB) (Gu et al., 2022) (https://microsoft.github.io/BLURB/leaderboard.html).

Relatively few works have been reported for the extraction of genetic variant impact. The Precision Medicine Track (track 4) in the latest BioCreative VI workshop (Islamaj Dogan et al., 2019) introduced a RE task for the extraction of experimentally verified protein-protein interaction (PPI) pairs affected by the presence of a genetic mutation. Some works for the relation extraction task in the BioCreative VI precision medicine task include an SVM classifier to identify protein-protein interactions that are impacted by mutations (Chen et al., 2017) and a convolutional neural network to predict the mutational impact on PPI relations (Tran and Kavuluru, 2017). Yet these works are limited to extraction of impact on a single event type (PPIs).

In the following sections, we first discuss the implementation of the eMIND system, specifically the development of the BERT model to extract the impact of mutations and the additional processing required for integration of the text-mined results in the additional bibliography of UniProt’s protein entry. We also describe an evaluation of eMIND’s impact extraction along with an analysis of the evaluation results.

## 3 Materials and Methods

### 3.1 Overall Approach

Figure 2 shows the computational components used in our pipeline to predict whether there is an impact of a genetic variant. Figure 2 depicts the workflow for the training of the BERT Relation Extraction (RE) classification model and the extraction of mutation impact relations on new abstracts. This involves the creation of the training set and fine-tuning the BERT model, which is discussed in sections 3.3.1 and 3.3.2. Next the trained BERT model on new test instances for impact relation prediction to extract the effector argument containing the mutation and the affected entity, which is either impact on protein molecular level or on biological processes. Test instances are created from new abstracts by splitting the abstract into sentences, parsing the sentence and detecting named entities (mutation and impact entities), which is discussed in sections 3.2 and 3.4. Details of the application of the trained BERT model on new test instances can be found in section 3.3.3. In the following subsections, we describe the components in further detail.

**Figure 2:**
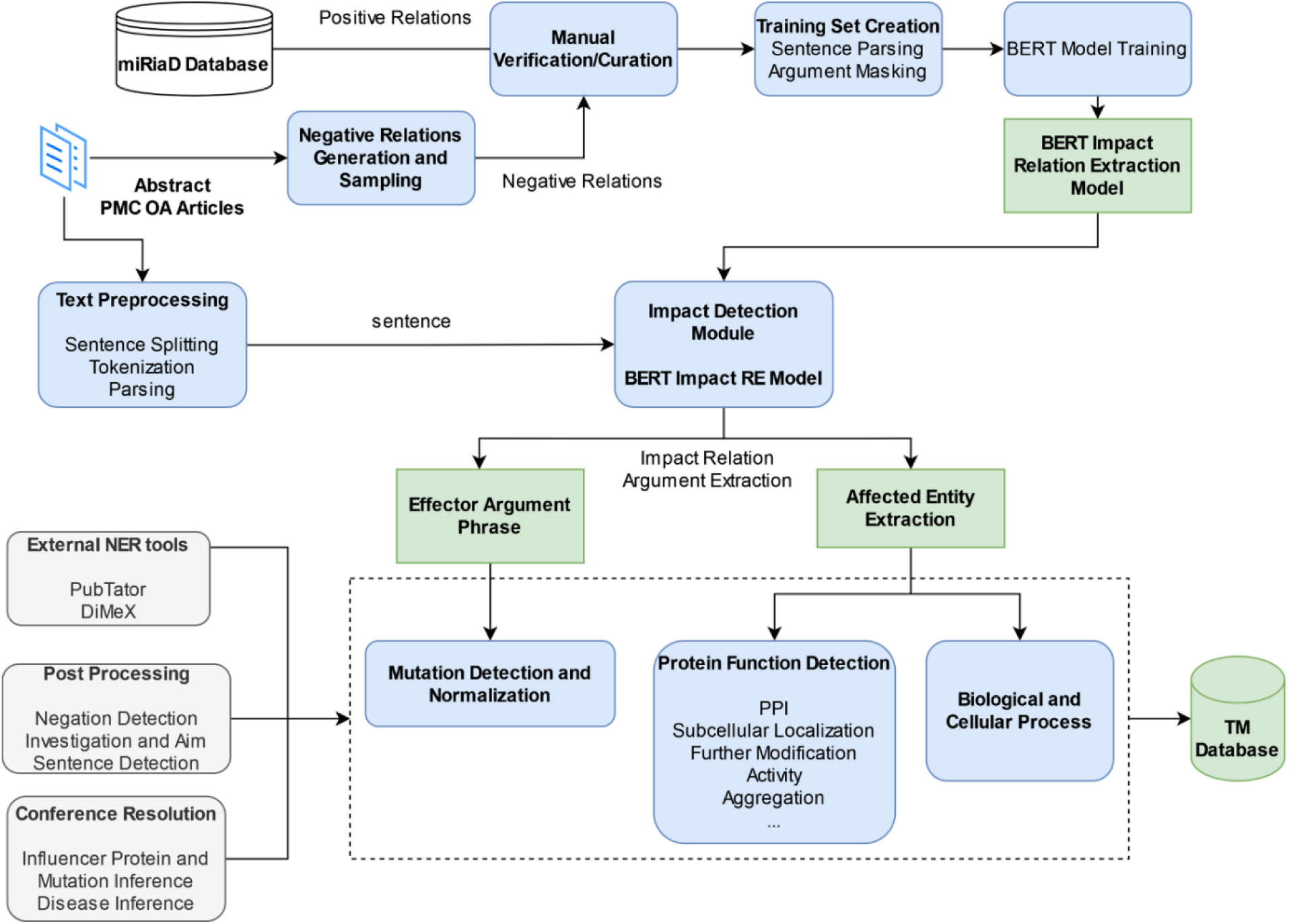
eMIND pipeline.

### 3.2 Preprocessing and Named Entity Recognition

We have processed and stored the entire Medline (as of January 2022) in a MongoDB database. Given an abstract, we first tokenize and split the text into sentences using the Stanford CoreNLP toolkit (Manning et al., 2014). We then use the Charniak-Johnson parser (Charniak, 2000; Charniak and Johnson, 2005) with David McClosky’s adaptation to the biomedical domain (Mcclosky, 2010) to obtain constituency parse trees for each sentence. Next, we use the Stanford conversion tool (de Marneffe et al., 2014; Manning et al., 2014) to convert the parse tree into syntactic dependencies, which provides a representation of grammatical relations between words in a sentence. We use these syntactic dependencies to extract the mutation and impacted entity phrases, which are typically Noun Phrases (NP).

Consider the sentence in Example 1 and its syntactic dependencies using the Universal Dependency notation (de Marneffe et al., 2021) (represented as a graph) depicted in Figure 3 below. Since the mutation and impacted phrases are typically Noun Phrases (NPs), we look at the outgoing edges from the nouns (called head nouns) in the syntactic dependency graph to obtain the corresponding NPs. For example, we follow the “nmod:of” and “compound” edges (indicating noun modification relations) from the head noun “mutation” to extract the mutation NP “*G206D mutation of presenilin-1”*. Similarly by following the “compound” edge we can extract the impacted entity NP “*Pen2 interaction”* headed by the noun “interaction”

**Figure 3:**
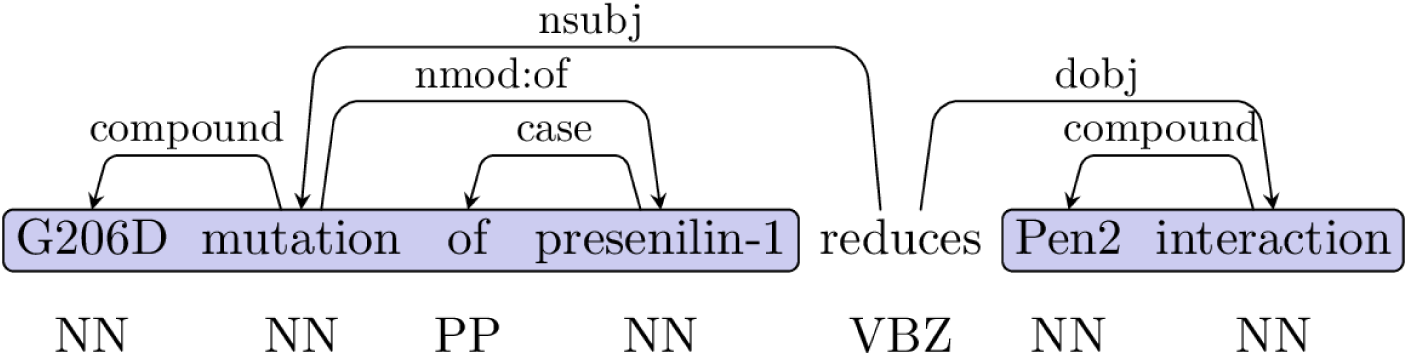
Syntactic Dependency Graph for Example 1. The mutation and impacted entity NP are highlighted in blue. Annotations (e.g. NN: singular noun, IN: preposition, VBZ: present tense verb) directly below the text correspond to the part-of-speech (POS) tags of each word in the sentence. The edges above the POS tags represent the syntactic dependency graph using the Universal Dependency notation. The syntactic dependency graph provides a representation of grammatical relations between words in a sentence. For example, the syntactic dependency edge from “reduces” and “mutation” is “nsubj” indicating mutation is the nominal subject of the verb “reduces”, while the edge “compound” from “interaction” to “Pen2” indicates a noun modification syntactic relation.

*Example 1: “G206D mutation of presenilin-1 reduces Pen2 interaction”* [PMID: 25394380]

Next, we use PubTator (Wei et al., 2013b, 2019), which is a precompiled resource that provides information about mentions of different types of named entities. In our context, we are specifically interested in the detection and normalization of the following types of entities: genes/proteins, diseases, and mutations. PubTator normalizes gene/protein mentions to NCBI gene identifiers, which are mapped to UniProt accessions using the ID mapping table downloaded from UniProt’s FTP site (https://www.uniprot.org/downloads). Similarly, the disease mentions are recognized and normalized by PubTator to MESH identifiers, which are then mapped to Disease Ontology ids (DOID). PubTator uses tmVAR (Wei et al., 2013a, 2018) to recognize mutations and normalize them to dbSNP identifiers when possible. In addition to the entities detected by PubTator, we do additional processing to identify terms of these types (genes, mutations and diseases) that are important for the impact extraction. For mutation terms, we also mark words such as “mutation”, “variant” and “polymorphism” since sentences that specify the impact of a mutation might refer to it by using phrases such as “this mutation”. For gene/protein detection we recognize several mentions of proteoforms that are not detected by PubTator and are relevant for AD/ADRD (such as, Abeta43 and APOE4). Thus, in a sentence such as the one in Example 2, we need to know that APOE4 is linked to AD to capture information about relevant impact.

*Example 2: “In addition, apoE4 did not bind beta/A4 peptide at pH < 6.6, whereas apoE3 bound beta/A4 peptide from pH 7.6 to 4.6”* (Strittmatter et al., 1993)

While PubTator links both APOE3 and APOE4 to APOE gene, the distinction of the specific APOE alleles is needed to properly extract the impact information. We have collected a set of genes with alleles and proteoforms that are frequent in the Alzheimer’s disease literature and have written patterns so that the alleles/proteoforms can be detected and from them, the gene/protein name can be identified and normalized. Finally, we have created a short list of names related to Alzheimer’s and other neurodegenerative diseases that are missed by PubTator. This dictionary is used to match text in the abstracts/sentences for detection purposes.

Note, we detect Noun Phrases (NP) that refer to the mutation, such as “v333D”, “this mutation”, or “the v33D variant”. In these cases, the word (noun), which heads the NP is either a specific mutation mention (e.g., “<u>v33D</u>”) or general mutation words such as “mutation”, “mutant”, “variant”, “polymorphism”, or “snp” (e.g., “this <u>mutation</u>”, “the v33D <u>variant</u>”). Additionally, since our BERT model is entity agnostic and extracts impact relations between heads of argument NPs, it can extract impact relation from NPs such as “carriers of v33d mutation”. Here the head noun (“carrier”) is neither a specific mutation nor a general mutation term but is of interest for further mutation impact processing due to the attachment of the mutational phrase (“v33d mutation”) to “carriers”. In these cases, we detect that an NP argument of an impact relation extracted by BERT contains a mutational phrase then later we will infer that this impact relation was the effect of the mutation.

There are other types of entities that need to be detected. These are relevant for determining the objects that are impacted by the mutation. These will be discussed in Section 3.4 below.

### 3.3 Impact Relation Extraction

#### 3.3.1 BERT Architecture

We use BERT (Devlin et al., 2019) for detecting the impact relation. There are two parts to developing a BERT model for any task. In the first stage, i.e., the pre-training stage, BERT learns general language representation for both the words and the text sequence through two unsupervised tasks: masked language model (MLM) and next sentence prediction (NSP). This stage is task neutral and differs from the second stage which learns how to make predictions for a specific task. In the use of BERT for relation extraction, it is almost universal to treat it as a binary *classification* problem. That is, in the present context, given an instance, a prediction of true (or 1) indicates that the model predicts that the instance contains an impact relation. To treat it as a classification problem, a piece of text (sentence in our case) is provided with two terms marked. The model predicts whether the text specifies an impact relation between the two entities given by the marked entities.

As is typical in the use of BERT for relation task, a sentence is provided as input after tokenization. A tokenizer such as SentencePiece (Kudo and Richardson, 2018) is used to break the sentence into subword tokens which are fed in the order they appear in the sentence. To indicate the two marked terms, we first mask the terms by the word “ENTITY” in the input. For classification purposes, the classification signal (CLS) in the last layer is used for prediction. A softmax layer on the output of BERT model is used for classification. During the fine-tuning of the model for classification, we use learning rate of 2e-5, batch size of 32, training epoch of 5 and max sequence length of 128.

#### 3.3.2 Creation of Training Dataset

A major hypothesis we are testing is that the structure and words used in statements that describe the impact of one entity on another entity or event/process are not specific to the two objects under consideration. This hypothesis was put to a test here since we created the training set based on the impact of microRNAs (also called miRNAs or miRs) to see if the text indicating the impact generalized to the impact of variants. We did this because of the need to create a large enough corpus for the training of a BERT model. miRiaD (Gupta et al., 2016) extracts relations from sentences in Medline abstracts that connect miRNAs and disease-related terms. Thus, miRiaD has a large amount of sentences that describe impact (of miRNA). Once we mask the miRNA mentions and disease-related terms from the sentences extracted by miRiaD, we hypothesize that the resulting instance should be structurally and lexically similar to text mentioning impact of a mutation. Consider the sentence captured by miRiaD “Expression of miR-140 regulates cell proliferation” for which a CAIR relation was detected by miRiAD of regulation between the arguments, “Expression of miR-140” and “cell proliferation”. For this positive instance a masked instance is created by replacing the heads of the arguments: “ENTITY of miR-140 regulates cell ENTITY”. Masking the head of arguments of CAIR relations from miRiAD will enable BERT to learn CAIR relations between head nouns of NP. Another hypothesis is that the same syntactic structure, learned through the language model of BERT for impact through CAIR relation can generalize to other domain. For example, in our case of capturing impact of mutations, a similar impact through “regulation” would be “V97L <u>mutation</u> increased the <u>levels</u> of Abeta42” (head nouns underlined). Using the miRiaD dataset we generated 8,245 positive and 11,496 negative relations for BERT fine-tuning. The negative relations were generated by performing a random sampling of NP pairs for abstracts in miRiaD database, where there is no CAIR relation detected by the miRiaD system.

#### 3.3.3 Application of the Model

Once the BERT model has been learned, it is then applied on instances to predict whether or not they specify an impact of a mutation. Our pipeline for the process of applying the BERT model is as follows. First, an abstract or a set of abstracts is chosen and based on the PMID of the abstract, the text is retrieved from our local MongoDB-based database. As mentioned earlier, the locally-stored abstracts are already split into sentences and parsed with the noun phrases noted. Simultaneously, we use the PubTator output (also stored locally) to identify the spans of text that correspond to named entities of types: gene/proteins, disease and mutations. Additionally, words such as mutation, polymorphism, and variant are also detected.

No lexical information regarding the entities is available during training because of masking of the head nouns. Thus, the learned BERT model will be unaware, for instance, whether the masked entity is a miRNA mention or anything else. Under the hypothesis that the structural or grammatical information will be useful, we consider the heads of noun phrases since they will be grammatically relevant arguments of impact. Since we learn entity-agnostic impact, it is important to note that these terms and named entity mentions may or may not be the terms that are marked. Because the training instance had the head of NPs marked as entities, we detect the NPs, base noun phrases and heads based on the parse of the sentence and then accordingly ensure that the head is masked as ENTITY.

Each sentence from a given abstract is considered and can give rise to 0 or more instances. An instance is created when a sentence has two masked entities, exactly one of which represents a mutation. Each instance is then fed to the BERT-based impact model which predicts whether or not the instance contains an impact relation between the pair of entities.

### 3.4 Impact Entity Recognition

As previously mentioned, one of the arguments of a positive relation predicted by the trained BERT model should be an impacted entity of interest (affected entity). The impacted entity phrases are detected and typed using a set of patterns and a list of trigger words. Recall, the impact phrases are typical Noun Phrases (NPs), which are the second argument of the impact relation. Our approach to detect impact entities is driven by the head word of the given NP and its attachments (determined through the dependency graph). Thus, given a sentence text, we first detect all NPs and associated head nouns (and their dependencies) and then apply a set of patterns based on the head noun to type the NP to the appropriate impacted phrase type. Our detection of NPs and associated head noun is based on syntactic dependencies as previously discussed in Section 3.2.

We follow outgoing edges from a noun to extract the NP headed by the noun. Thus, based on the syntactic dependency graph in Figure 3, for the noun “interaction”, we can extract the NP “Pen2 interaction” and thus this NP can be typed as “Protein Interaction” due to being headed by the noun “interaction”. Broadly the mutation impact phrases include among others, biological process, protein processing, protein abundance, and protein interaction.

#### 3.4.1 Protein-level Impact

Recall that the entities that are considered in this work are entities that are always NPs. Our patterns are mostly driven by first focusing on the head word of the potential impact entity NP. However, many times the head words are not sufficient, and we need to consider some context, i.e., the modifiers of these head words. In Example 2, consider the NP headed by the word “interaction”. We can only tag such NP as a protein-related impacted entity (protein interaction in our case) if the interaction involves a protein (e.g., “Pen2 interaction”) as opposed to other type of molecule (e.g., phospholipid interaction). Note that the impact can be on the variant itself or target a different protein. In Example 3, Psen1 mutations impact activity (loss of function) of Psen1, but also affects the abundance of Abeta (a target of Psen1).

*Example 3: These results support our previous conclusions that the L435F and C410Y mutations cause loss of Presenilin function and gamma-secretase activity, including impaired Abeta production in the brain* (Xia et al., 2016)

The detected protein-related impact is normalized to the closest term in the VariO Ontology (Vihinen, 2014). We briefly discuss each of these impact types below.

##### Effect on protein processing (VariO:0109)

One relatively common impact of a variant in AD/ADRDs is on protein cleavage and secretion. For example, any mutation affecting the cleavage of APP (including changes of Abeta proteoforms ratio) or shedding of TREM2 are examples under this impact type. In these cases, our detection is usually limited to looking for the head words such as “secretion” or “cleavage” and ensuring that a protein modifies this head word (e.g.: “Abeta secretion” or “secretion of Abeta”).

*Example 4: “Here we show that the Arctic mutation favors proamyloidogenic APP processing by increased beta-secretase cleavage, as demonstrated by altered levels of N- and C-terminal APP fragments.”* (Sahlin et al., 2007)

##### Effect on protein interaction (VariO:0058)

There are two types of interactions that we consider under this impact type. The first involves interaction between proteins, and thus involves looking for head words such as ’association’, ’binding’, ’recruitment’, ’interaction’ and a protein modifying the head word (e.g. “PS1-Pen2 interaction”). The second case involves the formation of a complex. Based on our study we noticed that many of NPs related to this type are headed with words such as “dimerization”, “trimerization” and “oligomerization”. Other times, the head word refers to the complex formation and, in such cases, the head words are “assembly” or “formation”. However, in these cases, the head word is not clearly indicative of a complex formation and hence we need to see the modifiers of these head words to be certain. These include dimer*, trimer*, oligomer* (e.g., “dimer formation”), as well as terms such as complex, channel or even a name of a complex (“assembly of FGF14:Nav1.6 channel complex”).

*Example 5: “Finally, our assay revealed that the Alzheimer’s disease-protective ’Icelandic’ mutation greatly attenuates APP-BACE-1 interactions, suggesting a mechanistic basis for protection.”* (Das et al., 2016, 1)[PMID: 26642089]

##### Effect on protein abundance (VariO:0052)

In this case the impact of variation is on the abundance/expression level of a protein. Some straightforward head words for this impact type are “level” and “expression”, once detected we check the presence of a protein as modifier of these head words. Additionally, we also consider terms such as transcription, production, proportion and ratio. The last two words (proportion and ratio) are often relevant in the context of AD/ADRDs because they might represent the ratio of peptides. Here we have to ensure that a word containing peptides separated by “-” or “/” modifies the head word.

*Example 6: “Overexpression of the p.D50_L51ins14 TYROBP mutant led to a profound reduction of TREM2 expression, a well-established risk factor for AD”* (Pottier et al., 2016)

##### Effect on protein post-translational modification (VariO:0107)

n these cases, we look for head words indicating the modification of proteins, such as “phosphorylation”, “glycosylation”, “ubiquitination”, “nitrosylation”, “methylation”, “acetylation”, “neddylation”, “sumoylation” and “lipidation” (e.g. “tau phosphorylation”, “Ser/Thr phosphorylation”). Note, since the presence of these head words is sufficient to indicate protein modification and hence we do not check the presence of the protein as a modifier to the head word. In some cases of protein modification impacted phrase, the above triggers might appear as a modifier (“phosphorylated”, “glycosylated”) to a head word, which can be a protein or a domain. Thus, additionally, we look for verb-forms of the words listed above modifying a protein/site head word.

*Example 7: “A single Y202F mutation in the Mint1 N-terminus inhibits C-Src binding and tyrosine phosphorylation.*” (Dunning et al., 2016)

##### Effect on protein activity (VariO:0053)

The detection involves looking for the head word “activity” and ensuring a protein modifies it. Certain modifiers such as the word “catalytic” or words ending with “ase” indicative of protein activity (e.g. “catalytic activity”, “secretase activity”) are also considered.

*Example 8: “Moreover, we found that the expression of the N141I PS2 FAD gene strongly promoted (>2-fold) glycogen synthase kinase (GSK)-3beta activity coincidental with a reduction in the level of beta-catenin translocated from the cytoplasmic to the nuclear compartment.”* (Qin et al., 2006)

##### Effect on protein aggregation (VariO:0038)

In these cases, we detected impacted entities indicating protein aggregation tendency. The detection of aggregation involves looking for head words such as “aggregation”. In addition to this, we look for terms indicative of protein aggregation in the context of AD. This involves looking for the head word “accumulation” with the presence of a modified protein describing protein aggregation. Additionally, some special head words indicative of protein aggregation include “deposit”, “plaque” (e.g. “fibrillar Abeta deposits”, “plaques of N- and C-truncated Aβ”).

*Example 9: “This study demonstrates that the Arctic APP mutation is sufficient to cause amyloid deposition and cognitive dysfunction…”* (Rönnbäck et al., 2011)

##### Effect on protein structure (VariO:0064)

In these cases, we detect impacted entities indicating protein structure in an effort to extract the impact of variants on protein structure and in such cases, the entity is normalized to the VariO term: “Effect on protein structure (VariO:0064)”. Trying to identify impacted entity phrases, which describe protein structure involves NP that mention words like “β-barrel”, “β-turn”, “alpha-helix”, “loop” etc. However, such NPs are typically headed by words like “structure” or “content” (e.g. “alpha-helix structure”, “β-turn content”). Additional set of head words that also indicate the structure of a protein include words ending with “mer” (e.g. “trimer”, “dimer”) with modifiers such as “loop”, “barrel” and “turn”.

*Example 10: “The L392V mutation of presenilin 1 associated with autosomal dominant early-onset Alzheimer’s disease alters the secondary structure of the hydrophilic loop.”* (Gantier et al., 1999)

#### 3.4.2 Biological Process

We follow the approach to detect biological processes that we had developed in emiRIT (Roychowdhury et al., 2021), a text-mining-based resource for microRNA information. A phrase is mapped to biological processes using a dictionary-based matching technique to obtain the closest match. The dictionary was created using a combination of terms and their synonyms from the cellular processes of the biological process branch of gene ontology (GO) (GO Consortium, 2021). We additionally augmented the dictionary derived from GO with terms mined from the text (PubMed abstracts) using patterns in text stating a collection process and member processes. For example, on encountering text such as “cellular processes such as migration, invasion and cell death”, we extract the three listed process terms. Such commonly appearing member process terms were mined from large amounts of Medline abstracts to create a comprehensive process dictionary. The detected biological process phrases are normalized to the GO identifier when possible.

### 3.5 Additional Processing

After the impact relation, mutation, and impact phrase typing has been completed we perform some post-processing to extract and normalize the associated disease and protein entities.

To detect disease entities, we first create a dictionary of diseases by combining PubTator’s disease output with a manually curated dictionary of neurodegenerative diseases. The in-house neurodegenerative disease dictionaries are generated by experts (biologists, curators) through manual review of neurodegenerative disease-related abstracts. Tags are included to mark subtypes of Diseases (e.g. EOAD, early-onset Alzheimer’s Disease). Diseases in the in-house dictionary were mapped to corresponding Disease Ontology Identifiers (DOID) entries. Based on this dictionary, we first look for the disease mentioned in the extracted evidence sentence for mutation impact. However, the disease might not be mentioned in the extracted mutation impact sentence, and thus we have to infer the disease from somewhere else in the abstract, following the order: abstract title, first sentence. Additionally, we look at certain “study” type of sentences in the abstract to detect the disease. These “study” sentences typically mention the experiment being conducted and might mention the disease central to the study. The detection of “study” sentence follows the approach of detecting “Patient context sentence” described in our work, DiMeX of associating mutation to diseases (Mahmood et al., 2016). We link the gene/protein of the variant detected by PubTator using the variant mutation to the gene association module of DiMeX, which are further normalized to UniProt Accession identifiers. Additionally, an in-house developed tool for Abstract sectioning, which is based on the work by (Karabulut and Vijay-Shanker, 2022) was applied so each evidence sentence extracted is assigned a “flag” indicating, from which section (“Introduction”, “Background”, “Method”, “Results”, “Discussion”, “Conclusion”) in the abstract the impact sentence was extracted.

### 3.6 Producing Output for UniProt

The development of eMIND was motivated by the general need of scaling up curation. We believe the contribution of publications with structured annotations to the computationally mapped bibliography section in UniProt entries will greatly expand the information available for users to explore, and could help curators to find publications for further curation. Thus, the output of eMIND needs to be prepared in a tab-delimited format to be included in the UniProt bibliography pipeline. The format consists of the following information: UniProt accession|source|link|PMID|[Category]Annotation. In the case of eMIND, the category for the publications corresponds to the UniProt annotation topic covering variants. The annotation is structured with Variant:HGVS nomenclature. Impact:VariO or GO for process. Evidence sentence and abstract section where it was extracted. Here are two such examples:

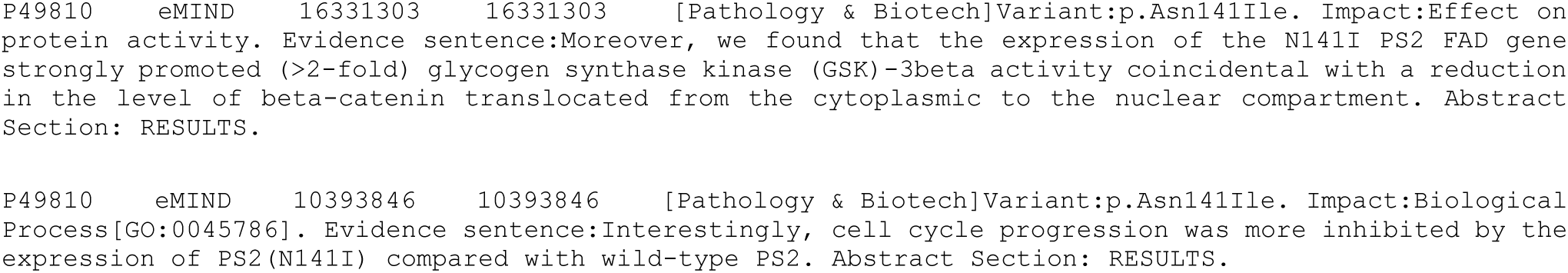

Figure 4 below shows a screenshot of the corresponding UniProt entry UniProtKB:P49810 (Presenilin-2) indicating how the annotations above appear in the publication section (partial snapshot).

**Figure 4:**
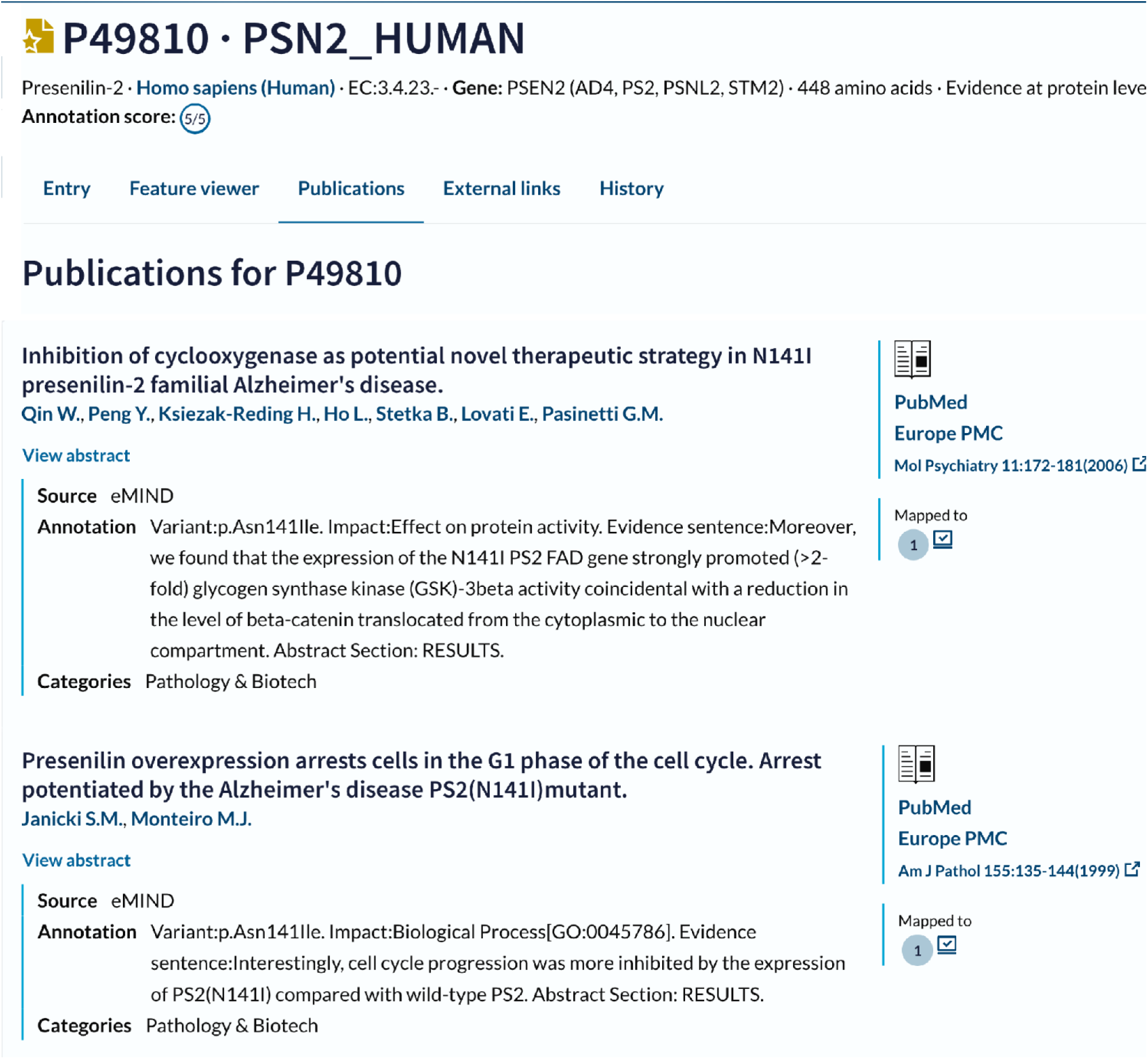
Sample eMIND’s text-mined output integration in UniProt for protein entry Presenilin-2.

As a pilot, a small sample of eMIND’s text-mined output inspected by the curator were included in the UniProt additional bibliography section. In the future, the UniProt additional bibliography for protein entries will be regularly updated based on running eMIND on the set of abstracts recommendations provided by a LitSuggest model (Allot et al., 2021), which will be trained using the abstracts currently integrated into UniProt.

### 3.7 Evaluation Dataset

#### Selection of the dataset

The test set was composed of 60 abstracts that were selected from three different sources to provide a variety of examples about the impact of protein mutations on AD/ADRDs.

1-UniProt: We downloaded the information from protein entries from the AD disease portal (Wang et al., 2021), UniProtKB Release 2021_04), which contains proteins that are candidates for AD (Breuza et al., 2020). From these, we selected 15 publications that constitute the evidence from existing UniProt annotations of natural variants involved in AD/ADRDs for 12 different proteins. Specifically, those containing a note describing impact after the mention of disease name. For example, from P10636, VAR_019660 corresponding to p.Arg5His, has annotation: (in FTD; reduces the ability of tau to promote microtubule assembly and promotes fibril formation in vitro; dbSNP:rs63750959) evidence=”ECO:0000269|PubMed:11921059.” We reviewed examples and selected those where the impact information was present in the abstract.

2-AD computational bibliography set: UniProt computationally mapped bibliography contains different publications collected from a variety of sources. One source is from our literature mining preliminary method based on EDG and a small number of handcrafted rules about variants with impact on AD. We randomly selected 35 abstracts about variants with impact on the protein-related aspects of interest.

3-LitSuggest (Allot et al., 2021): We collected an additional set of abstracts directly from the literature using LitSuggest. The set of articles described in 2 was used as the positive set to train the LitSuggest classifier with impact in ’Alzheimer’, with options include_mutation: true, include_gene: true. The negative set was automatically selected by LitSuggest. After training the classifier, the new set of articles suggested were reviewed to indicate relevancy to impact, and based on this new annotation, the system was retrained on new relevant articles. From that set, the first 12 relevant publications were included in the set.

#### Annotation and Annotation framework

Two curators reviewed a number of examples on protein mutation impact on AD/ADRDs and created the annotation guidelines for the project. PubTator was used to pre-annotate entities of interest (genes, variants, disease, species). PMIDs in the evaluation exported from PubTator central in BioC format. TeamTat (Islamaj et al., 2020) was used as the collaborative framework for text annotation. Documents with annotations in BioC were imported into the system and annotated by two curators.

Figure 5 shows an example of an annotated document.

**Figure 5:**
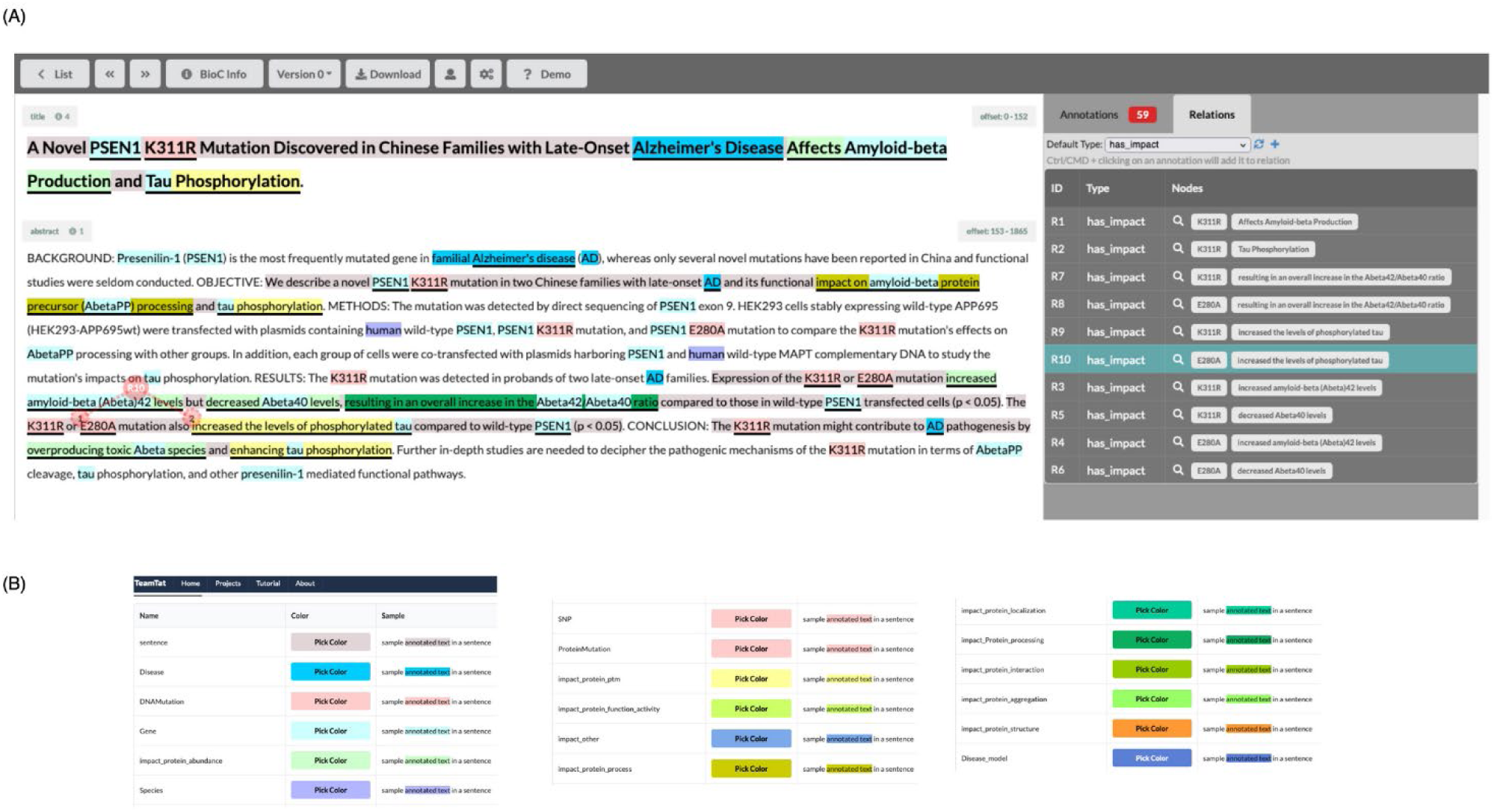
Evaluation corpus annotation interface. (A) Abstract with entities highlighted. The right panel shows the impact relations annotated. (B) Color-code for the entities annotated.

## 4 Results and Discussion

The evaluation set consisted of annotations of impact relations from 60 abstracts about AD/ADRDs, covering 25 gene targets involved in impact relations for a total of 69 unique variants (top 10 gene shown in Table 3). Figure 6 shows that the dataset covers all relations impact types of interest, with most annotations involving the effect of mutation on protein abundance.

**Figure 6:**
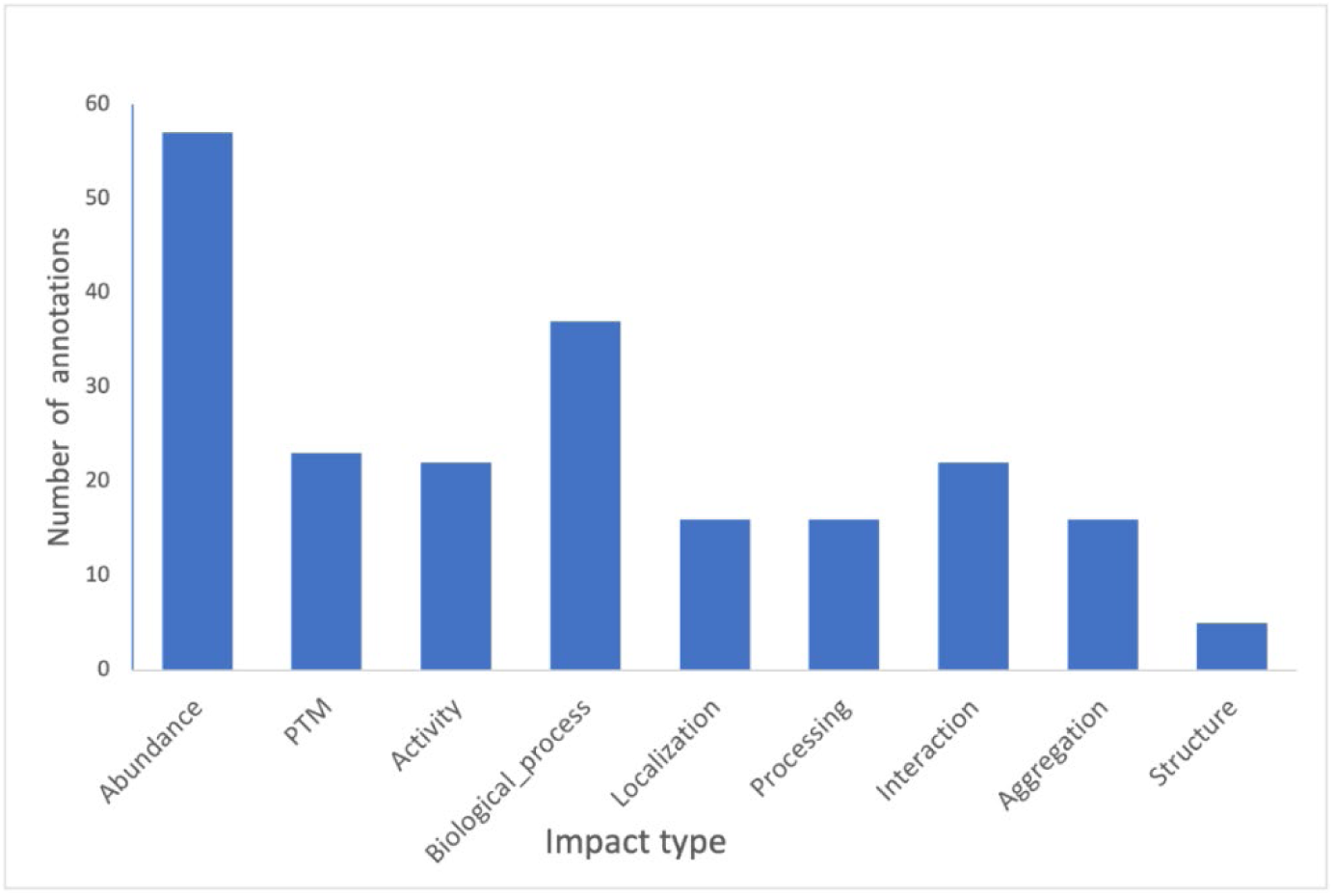
Distribution of annotations by impact types in the evaluation dataset.

**Table 3:** Top 10 Gene coverage and variant frequency in the dataset.

| Gene | Unique variants | Total # variants |
| --- | --- | --- |
| APP | 17 | 20 |
| PSEN1 | 11 | 13 |
| MAPT | 8 | 10 |
| TREM2 | 7 | 13 |
| SORL1 | 3 | 3 |
| BDNF | 2 | 2 |
| DNM1L | 2 | 2 |
| ADAM10 | 2 | 2 |
| APOE | 1 | 2 |
| CALHM1 | 1 | 2 |

The annotated instances were positive instances only and corresponded to sentences where there was a relevant impact of a mutation. The impacted entity and mutation phrases were marked as well. Altogether there were 163 positive instances. These instances/abstracts formed the basis for the dataset used in our evaluation. All 163 instances were chosen for the recall study. On the other hand, for the precision study, we started with 100 instances predicted to be positive by running our system on these abstracts. The evaluation for precision involved deciding how many of them were indeed positive and how many of them are false positives.

Using the evaluation set, we conducted two evaluations. First, since the domain experts’ annotations only included positive instances and no negative instances (and neither would it be appropriate in this case to assume non-positively annotated instances are negative), we used the positive instances to compute recall. That is, we ran our system on the abstracts that had been annotated and considered how many of the positive instances were predicted to be positives by the system and how many were not. The second evaluation considered the positive instances predicted by the system. We randomly chose 100 such instances and manually decided whether they were correctly predicted or not. Recall, the expert-annotated instances only included positive sentences and that an instance could still be positive even if it is not included among the expert-annotated instances. To evaluate the precision, we considered the same set of abstracts that were used in the recall study.

**Recall**: Of the 163 positive instance annotations, our system was able to correctly identify 136 instances. This gives us a recall of 0.84 percent. We note that 14 of 27 missed cases represented the situation where our system (possibly due to impact entity recognition) didn’t propose an instance to be submitted to the BERT impact predictor. Hence, clearly, this is a false negative for the system since this instance was not predicted as positive by the entire pipeline.

**Precision:** Of the 100 instances we examined, we found 94 were correct and that there were only 6 false positives. In fact, as discussed below in the analysis of errors, detection of impact relation was not an issue and the BERT model seems to hardly predict any false positives for the impact relation.

### Error Analysis

We conclude that the BERT impact model generalized well from its training data and that almost every false-negative error was due to the sentence not being similar to the impact instances in the training set. There were two main types of sentences that the BERT model failed to capture. In the first case, the impact (or association) of the mutation was specified by using a contrast or comparison. For example, *“… although the mutant protein is expressed, it is not secreted, …”* (Mukherjee et al., 2008). Annotation of the impact relation relies on the knowledge of the wild-type protein’s secretion status. As far as we know the training set didn’t include any such associations made because of the contrast or comparisons (other cases). The second type is exemplified by instances like *“Interestingly, the AD-associated PS1(L166P) variant revealed a partial loss of γ-secretase function, resulting in the decreased production of endogenous Aβ40 and an increased Aβ42/40 ratio.”* (Koch et al., 2012). In this example, we are focusing on the impact specified by “partial loss of gamma-secretase function”. There were other such examples where verbs such as “exhibited”, “displayed” and “showed” were used in place of “revealed” in the above sentence followed by the mention of some change (“loss” in this case). Again, the training examples did not have such examples as they were not detected by miRiaD.

The false positives of our system were not due to the detection of impact relation by the BERT model but for other reasons. Two of the errors were due to the fact one of the masked arguments presented to the BERT model was not a mutation phrase. The errors were due to the fact that our recognizer incorrectly considered G1254023X as a mutation mention whereas in actuality it is a chemical (metalloproteinase inhibitor). Other errors were due to incorrect detection of impact entity or their type. For example, in *“A non-synonymous polymorphism, rs2986017 (p.P86L), in the newly characterized calcium homeostasis modulator 1 (CALHM1) gene located in the Alzheimer dementia (AD) linkage region on 10q24.33, was reported to increase risk of AD, and affect calcium homeostasis and amyloid beta accumulation.”* (Sleegers et al., 2009), “(calcium) homeostasis” was recognized as a biological process, whereas in this sentence this is only part of a gene/protein name and hence shouldn’t even have been considered as an impact argument and masked.

## 5 Future Work

eMIND impact relation extraction module considers impacts conveyed through CAIR relations (Connections through Association, Involvement, and Regulation) as indicated earlier in Section 2.2. Based on the errors analysis i.e., impact sentences missed by eMIND on the evaluation set and further analysis on a development set, we noticed additional constructs that can convey impact of genetic variant. Table 4 below provides sample example sentences for three additional constructs we plan to incorporate into eMIND in the future. One such construct (row 1) involves cases that convey the impact relation through comparisons, where an alternate protein form (e.g., a mutated protein) is compared with normal/control (e.g., wild-type). Comparison is indicated with words such as more, less, increased, elevated, enhanced etc. Notice the normal form (“wild-type) is explicitly stated in the example sentence in row 1 but might be omitted and left unstated as in rows 2 and 3. In these cases, typically we have words such as “showed”, “displayed”, “exhibited” etc., (row 2) or phrases such as“leads to”, “resulted in” etc. (row 3) before the comparison words (“decreased”, “enhanced”). In these two cases, the comparison words indicate a Change of Value (COV) from another situation (the unstated normal case). In addition to incorporating the additional impact constructs noted here, we also plan to conduct a new evaluation to measure the performance of eMIND on extracting these impact constructs. Additionally, we plan to run eMIND on a large scale on all AD/ADRD-related Medline abstracts and disseminate the text-mined results through our tool, iTextMine (Ren et al., 2018).

**Table 4:** Additional impact relation constructs.

| Relation Construct | Example Sentence |
| --- | --- |
| **Comparison/Contrast** | <u>Compared with</u> its wild-type, NDPK-A (S120G) appears **<u>more</u>** effective in promoting neuroblastoma metastasis. |
| **Show-Change of Value (COV)** | Carriers of this 'Arctic' mutation <u>showed **<u>decreased</u>**</u> Abeta42 and Abeta40 levels in plasma. |
| **Leads_to-Change of Value (COV)** | .. the p.H157Y variant within the stalk region <u>leads to **<u>enhanced</u>**</u> shedding of TREM2. |

## 6 Conclusion

In this work, we presented the text-mining tool eMIND to extract the effect of mutations in the context of AD/ADRDs. Although eMIND is able to capture the effect of a mutation for both coding and non-coding variants, in this work we focused on the impact involving protein coding variants, given its specific application in associating publications and annotations to UniProt entries. For this purpose, we reviewed the type of impact at the protein level that UniProt captures (such as protein activity, aggregation, processing, modification, structure, interaction, involvement in biological process and abundance) and followed annotation guidelines on variants.

eMIND uses BERT, a transformer-based model to capture the impact between biomedical entities and processes. This in combination with a rule-based named entity detector to detect impacted mutation entities (protein activity, processing, etc.) enables us to capture the effect of mutation on the various protein properties. A study was conducted to evaluate eMIND’s approach to extract mutation impact relations and we obtained a recall of 0.84 and a precision of 0.94. A sample of eMIND’s text-mined output has been integrated into UniProt’s additional bibliography section of different protein entries. We aim to update the additional bibliography of UniProt’s entries with eMIND’s output by running eMIND on a weekly basis on abstracts suggested by LitSuggest. As indicated earlier, we plan to run eMIND on a large scale on all AD/ADRD-related Medline abstracts and disseminate the text-mined results through our tool, iTextMine (Ren et al., 2018).

## Conflict of Interest

The authors declare that the research was conducted in the absence of any commercial or financial relationships that could be construed as a potential conflict of interest.

## Author Contributions

SG developed the BERT-based version of eMIND and was involved in writing the manuscript first draft; XQ developed the first version of eMIND; QW was one of biocurators annotating the evaluation dataset; JC was involved in development of eMIND website; HH was involved in the integration of eMIND output into the additional bibliography pipeline in UniProt, CW, as one of the project PIs, contributed with original system idea and funding, KVS was involved in the design of the methods developed and contributed to the first draft, CNA contributed to original system idea, annotation of evaluation dataset and first draft. All authors have read and provided their feedback to this manuscript.

## Funding

This work has been funded by NIA supplement grant to UniProt 3U24HG007822-07S1, NIH/NHGRI: UniProt - Enhancing functional genomics data access for the Alzheimer’s Disease (AD) and dementia-related protein research communities.

## Acknowledgments

We would like to acknowledge the UniProt Consortium (https://www.uniprot.org/help/uniprot_staff).

## Data Availability Statement

The annotation guidelines and evaluation annotated corpus generated for this study can be found in the iProLINK [https://research.bioinformatics.udel.edu/iprolink/corpora.php].

## Notes

https://research.bioinformatics.udel.edu/itextmine/emind

https://research.bioinformatics.udel.edu/iprolink/corpora.php

### Competing Interest Statement

The authors have declared no competing interest.

